# Printed Droplet Microfluidics for on demand dispensing of picoliter droplets and cells

**DOI:** 10.1101/167163

**Authors:** Russell H. Cole, Shi-yang Tang, Christian A. Siltanen, Payam Shahi, Jesse Q. Zhang, Sean Poust, Zev J. Gartner, Adam R. Abate

## Abstract

Although the elementary unit of biology is the cell, high throughput methods for the microscale manipulation of cells and reagents are limited. The existing options are either slow, lack single cell specificity, or utilize fluid volumes out of scale with those of cells. Here, we present Printed Droplet Microfluidics, a technology to dispense picoliter droplets and cells with deterministic control. The core technology is a fluorescence-activated droplet sorter coupled to a specialized substrate that together act as a picoliter droplet and single cell printer, enabling high throughput generation of intricate arrays of droplets, cells, and microparticles. Printed Droplet Microfluidics provides a programmable and robust technology to construct arrays of defined cell and reagent combinations and to integrate multiple measurement modalities together in a single assay.

## Introduction

While the basic building block of life is the cell, humans comprise trillions of molecularly distinct cells whose interactions give rise to the functional properties of our bodies. A more detailed understanding of this network of interacting cells is required to reveal the etiology of human disease and to enable the regenerative medicines of the future, yet its complexity defies detailed study *in situ*. However, current techniques for cell manipulation, isolation, and combination are rudimentary and of limited value for high throughput studies, particularly when multiple cells of different type are required for the phenomena of interest.

Cell suspensions can be loaded into microfluidic droplets or wells to isolate large numbers of cells, yet the uncontrolled and statistical nature of cell loading leads to many volumes that are unusable because they are empty or contain undesired cellular contents. Deterministic dispensing to well plates with fluorescence-activated cell sorting is an efficient single cell handling technique, though depositing cells into microliter volumes, precludes many advantages of assays in small volumes, complicating the formation of direct cellular contacts, and diluting secreted molecules to non-physiological concentrations. Tackling problems with increased cellular complexity requires that biologically-meaningful experiments be carefully constructed and performed with massive reductions in reactor volume and significant increases in throughput.

In this paper, we address this fundamental limitation of current technologies using Printed Droplet Microfluidics (PDM), a new kind of microfluidics that prints droplets containing reagents and cells to defined arrays, where they can be deterministically combined and manipulated. The core of the approach is a fluorescence-activated droplet sorter (1) acting as an intelligent cell and droplet dispenser coupled with a motorized stage. Candidate droplets are cycled through the instrument and scanned to determine their contents, then printed to a substrate if a droplet contains the reagent or cell required for a given position in the array. In this way, droplet printing is made deterministic by intelligent sorting, allowing the construction of cell and reagent arrays of unprecedented intricacy. PDM combines the control and generality of microliter pipetting in well plates, the precise single cell dispensing of FACS, and the scalability and single cell sensitivity of droplet microfluidics.

## Results

### A microfluidic robot for droplets and single cell dispensing

Microliter pipetting has become a universal tool in biology due to its ability to deliver precise volumes of multiple reagents into well plate reactors for subsequent assays. These tools are easily reconfigurable using software, allowing the same instrument to be used in applications as diverse as studying cancer cells to engineering bacteria with unnatural properties(2, 3), and are now broadly used across the biomedical sciences.

Microliter fluid handlers, however, retain two major functional limitations: the microliter volumes required limit the number of reactions that can be performed; and the methods for dispensing liquids are not optimized to deliver precise numbers of individual particles or cells. To increase throughput, volume must scale down, which is the core concept of droplet microfluidics: a field that studies reactions in picoliter droplets six orders of magnitude smaller than fluid volumes typical in microtiter plates. By operating at the scale of individual cells, droplet microfluidics affords massive increases in speed and reaction number compared to microliter pipetting. Additionally, by combining droplet microfluidics with an in line fluorescence-activated droplet sorter, it becomes possible to insure that every droplet contains one and only one cell. However, droplet microfluidics remains an incomplete solution for picoliter fluid handling, even when combined with in line droplet sorters. Specifically, the process of forming, combining, sorting, and analyzing picoliter droplets requires custom microfluidic devices unique to each task. Therefore, each new task requires a new device and an expert to design, build, and operate it. To impact biology with better fluid handling, a new approach is required that achieves the throughput and scalability of droplet microfluidics, while retaining the flexibility and reconfigurability of microliter fluid handlers.

Several technologies have been developed for the controlled and flexible manipulation of microfluidic droplets. Digital microfluidics utilizes electrowetting to move, combine, and split nanoliter-scale droplets on a specialized substrates, and have been used to prepare samples for sequencing (4) and to probe inter-cellular signaling on the single cell level (5). A number of approaches have used open-ended microfluidic devices to dispense nL volumes to substrates(6, 7). Though these technologies enable droplet microfluidics in arrayed formats, the ability to manipulate single cells and rapidly switch reagents is lacking.

We have therefore developed a new microfluidics technique that further breaks through the constraints of conventional droplet microfluidics. Our method, PDM, miniaturizes array pipetting to the picoliter scale. The core of the technology is a microfluidic sorting-on-demand device that acts as a deterministic picoliter droplet dispenser on a motorized substrate (Fig. 1). The key attribute of this device is that it can select a specific droplet from a set of candidates and dispense it to a substrate (Fig. S1a, device schematic, Fig. S1b). It does this in a completely deterministic and programmable fashion, allowing the printing of intricately defined arrays of picoliter to nanoliter volumes based on a software encoded print file. Moreover, it is fast, selecting from a set of candidate droplets at up to kilohertz frequencies (8).

**Figure 1.**
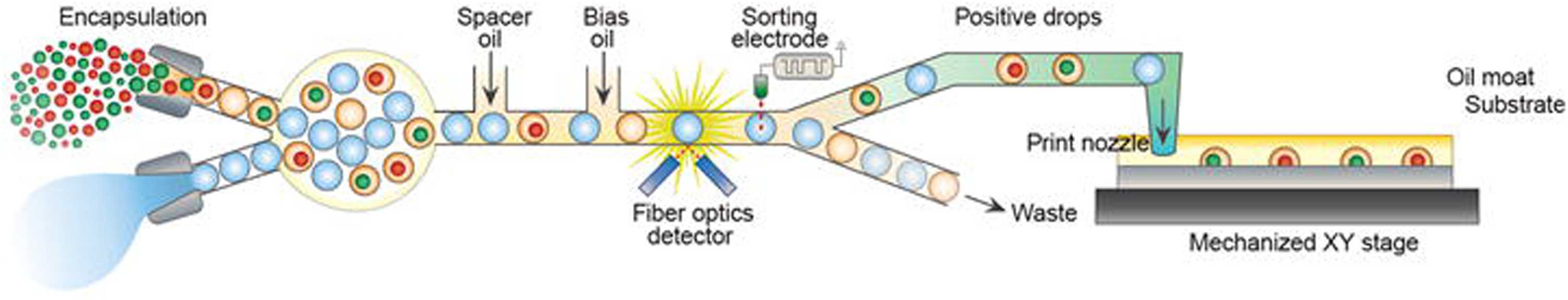
The printer consists of a sorter-based print head that selectively dispenses microfluidic droplets to a substrate under a cover of oil. The printing process is automated by coupling the print head to a motorized stage. Deterministic combinations of cells and reagents can be formed at each location through additive droplet printing.

To implement a microfluidic sorter as a droplet printer, we overcame several technical challenges. Foremost, droplet sorting requires oil as a carrier phase that must be displaced for the droplets to reach the substrate. To overcome the viscous and buoyancy forces that would slow or prevent deposition of droplets, we require a counter force to pull the droplets to the print locations. A simple and general technique for this is dielectrophoresis, commonly used to pull conductive droplets in oil for high speed sorting (1). To implement dielectrophoretic trapping, we construct a substrate consisting of an array of bipolar electrodes under a thin sheet of dielectric (Fig. S1c). When energized, the electrodes emit field extending above the substrate, causing droplets to settle into nanowells patterned on top of the dielectrophoretic traps where they are immobilized (Fig. S1d).

### Fundamental fluidic operations

Single droplet delivery is accomplished by positioning the outlet nozzle of the sorter above a dielectrophoretic target, dispensing a single droplet by activating a dielectrophoretic field in the print head to divert a droplet to the exit channel. The droplet is pulled into the trap minimum, displacing the oil as it reaches, and touches, the substrate (Fig. S2a). The substrate electrodes are covered by an insulating film to prevent current flow within an immobilized droplet.

A persistent challenge in droplet microfluidics is that it is difficult to add many reagents to pre-existing droplets. Current methods, such as coalescence (9) and picoinjection (10) are effective for a limited number of additions, and not well suited for the multitude of workflows that require reagent additions to occur at different times. In PDM addition can be accomplished by hovering over a printing position and dispensing additional droplets (Fig. S2b). Each dispensed droplet is pulled to the trap minimum, where it immediately electrocoalesces with the droplet already positioned there. The ease with which PDM can perform deterministic and programmable addition of reagents to preexisting droplets makes it possible to perform a suite of manipulations that are too impractical to reliably perform with flowing droplet microfluidics.

Droplet dispensing and addition are the basic operations for generating arrays with defined composition. Often, the next step in an experiment is to image the array and, in some cases, recover select droplets for further study, such as additional rounds of culture or sequencing. Recovery can be accomplished by simply vacuuming the droplet from the substrate, even with the electric field on (Fig. S2c). Recovery of mass numbers of droplets from a printed array via suction requires that a suction nozzle raster over the entire print substrate. With the current hardware, indiscriminate recovery of droplets from the array can be performed at ~4 Hz, and is essentially limited by the rate of stage movement. Recovery of contents from individual positions and placement of these fluid volumes into specific wells in a well plate is slower to allow for the transit of individual droplets to a unique external location and can be performed at ~0.1 Hz.

### Programmed printing of intricate droplet arrays

Microliter pipetting is used universally in the biomedical sciences due to its flexibility and the control with which it can combine multiple reagents. Picoliter dispensing technologies such as PDM must thus be able to create arrays of comparable definition and complexity to ensure similarly broad utility. Competing technologies for scaling-down reagent transfers rely on inkjet or acoustic dispensing (11), yet these technologies are used to construct experiments at microliter scales to prevent rapid droplet dehydration. Furthermore, constructing combinatorial experiments requires either additional printing channels, or a time-consuming reagent switchover step, a proposition that becomes onerous for highly combinatorial screens. By contrast, PDM switches elegantly between reagents contained in a mixed emulsion “ink” by dielectrophoretic droplet sorting, possible at kHz rates (8).

The current system print rate is related to three relevant delay times: the wait time for a desired droplet to enter the detection region of the print head, the droplet travel time from the print head sorting junction to the printing substrate, and the time for the stage to move from one nanowell array position to the next. For the system described here, droplets are typically generated or reinjected at 50-100 Hz, leading to wait times of 10-20 mS between uniform dye droplets and 0.2-0.4 S between droplets encapsulating single cells (at 1:20 limiting dilution). The travel time for a droplet to reach the substrate is ~0.1 S, and the mechanical stage can move between positions 400 μm apart in a 0.25 S. Our control algorithm enables droplets to be sorted with a fixed time delay, so that the 0.1 S of droplet travel occurs during stage movement, and does not limit the maximum 4 Hz print rate of the system when droplets delay times are short. Because multiple droplets can be sorted and en route to the substrate at a given time, printing multiple dye droplets to a position requires that the print nozzle hover over the print substrate for the additional sorting time associated with these droplets. For cell experiments in this work, we chose extremely selective gates to eliminate the possibility of sorting cell doublets, which in turn limited the availability of desired cells to ~1 Hz. Although beyond the scope of the proof of concept experiments demonstrated in this work, increasing printing throughput requires several straightforward system modifications. Our lab has demonstrated dielectrophoretic droplet sorting at rates of up to 30 kHz (8), so detecting and sorting droplets at 10-100X greater frequencies is readily achievable by increasing oil and reagent flows. Similarly, the time scale for droplet travel to the substrate is reducible by 10-100X through increased flow and decreased channel dimensions. The use of a faster piezo stage will enable the print head to move between 400 μm array positions at ~ 20 Hz, and address 10,000 position arrays in ~ 10 minutes.

Sorting-on-demand requires that the contents of each candidate droplet be identifiable via in-flow fluorescence in the microfluidic sorter. Since most reagents of interest are not fluorescent, we therefore include “fluorescence barcodes” in the droplets. For example, 100 different reagent solutions can be labeled by mixing two dyes at ten concentrations. Such combinatorial labeling has been demonstrated using lanthanide nanophosphors for over 1000 unique spectral barcodes, indicating that reagent labeling is not a meaningful limitation (12).

To demonstrate PDM’s ability to construct intricate arrays, we generate arrays of defined composition from two solutions of fluorescent dye, green (FITC) and red (TRITC) (Fig. 2). The dye solutions are injected via syringe pumps and encapsulated into droplets in a double T-junction upstream of the sorter (Fig. 2a, *inset*). The device forms droplets of both solutions in an alternating sequence, which is used to print the desired combination via sorting-on-demand. The droplets are scanned for fluorescence using integrated fiber optics, appearing as two tight clusters on a red versus green scatter plot (Fig. 2a). These clusters define “red” or “green” population gates, allowing the instrument to print them in a deterministic fashion following the print file.

**Figure 2.**
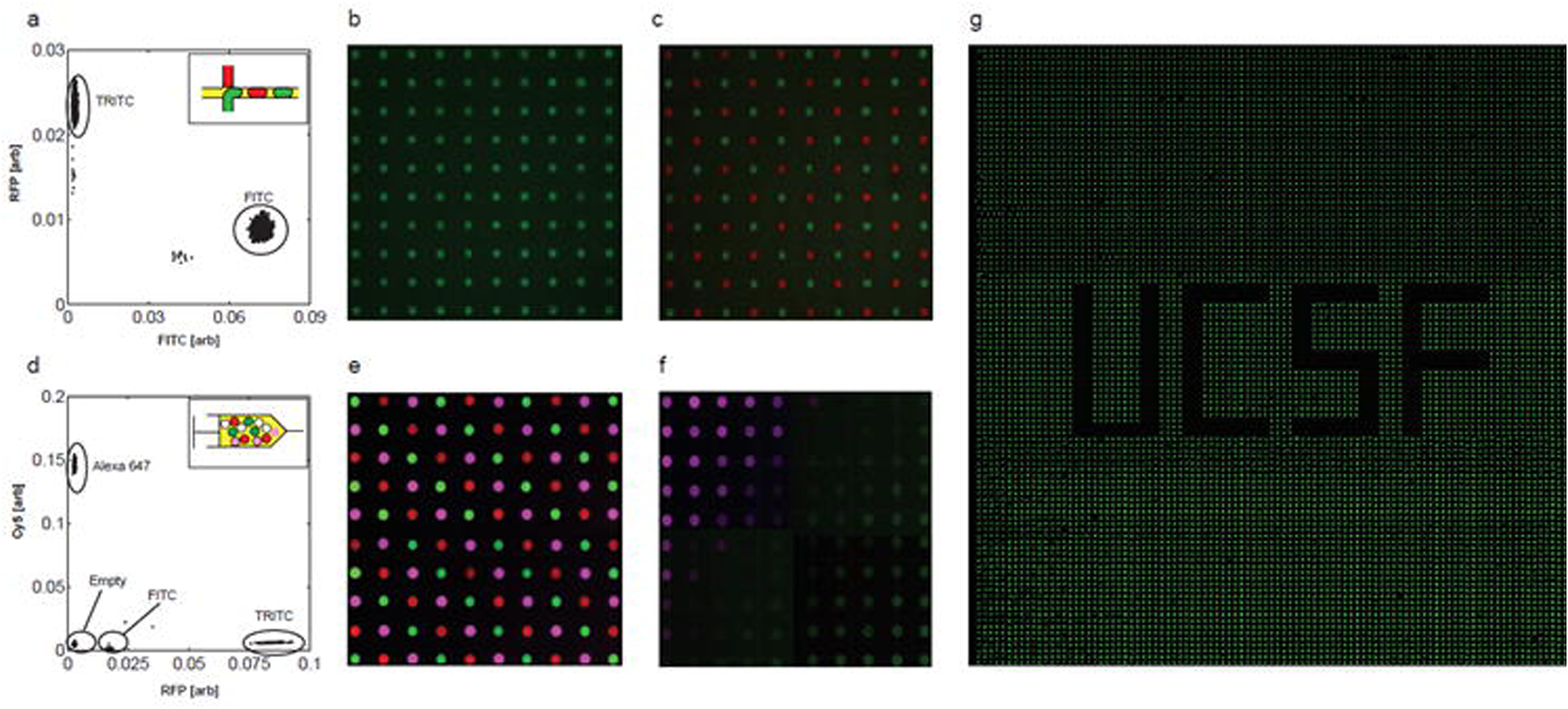
Fluorescent dye printing. a) Flow cytometry data for print head utilizing a dual drop maker to produce emulsions on chip. b) 100 position array printed with 4 droplets of FITC at each position from the emulsion shown in A. c) 100 position array with alternating positions printed with 4 droplets of either FITC or TRITC. d) Flow cytometry data from a reinjected emulsion containing 4 dyed droplet types. (empty, FITC, TRITC, and Cy5 (Alexa 647)) e) 100 position array printed with alternating positions containing 4 droplets of FITC, TRITC, and Cy5. f) position well gradient array printed with 8 droplets containing combinations of FITC, Cy5, and empty droplets. g) Construction of a binary image by printing FITC droplets to a 10,000 position array.

The simplest printing pattern is an array of identical droplets with identical contents. To print this array, we program 4 green droplets to be printed into every array position in the print file, and print the file. The instrument follows the program, sorting 4 green droplets into every spot on the array, yielding a uniform population of 100 green droplets, each composed of 4 combined droplets, and each immobilized in a nanowell structure located above a dielectrophoretic trap (Fig. 2b). Uniform sets of microscale reactors like this, however, can be created with many methods, including common microfluidic droplet generation, passive filling of nanoliter wells, or slip-chip systems (13–15). A more complex pattern that is impossible to build with these methods is alternating red and green droplets. To construct this array, we program the print file appropriately, and print it. The instrument sorts the droplets on demand in accordance with the file, generating the desired alternating array (Fig. 2c). In this way, changing the experiment performed by the instrument requires only re-writing the print file, rather than changing hardware – a departure from conventional flowing droplet microfluidics.

Encapsulating reagents on the device using a double T-junction is simple and allows printing of two solutions. However, this becomes impractical when many reagents must be printed, because this would require many feed lines connected to the tiny ~1 cm^3^ sorting device. A better approach is to emulsify all solutions, combine them in a mixed emulsion, and reinject this mixed emulsion “ink” through a single line. To illustrate this, we generate a mixed emulsion of four droplet types, labeling them with unique combinations of FITC, TRITC, Cy5, and Cascade Blue (CB) dyes. The detection scatter plot has four clusters, one for each droplet type (Fig. 2d). We use these to define color gates and load a print file with a pattern that prints 4 droplets of 3 different types to each position in a organized mixed color array (Fig. 2e).

Arrays like these illustrate the potential for using PDM to design and execute totally independent reactions at each spot on an array. Often, however, the objective will be to scan a parameter space that depends on the relative concentrations of multiple reagents. This can be accomplished by systematically varying combinations across the array (Fig. 2f). A major advantage of PDM over flowing droplets is that grid locations relate the composition of every droplet in the printed array; the array is generated in a deterministic fashion using a print file that contains all information about what was combined at each spot. While in flowing droplet microfluidics it is possible to generate varying concentrations of reagents using oscillating flows or random droplet merger (16, 17), it is far more difficult to determine what was combined in each droplet afterwards, since they are not neatly stored on an array.

Another key advantage of PDM is the ability to perform experiments at high throughput. To demonstrate this, we use green droplets to print an image file to a 10,000-position array (Fig. 2g, higher resolution Fig. S3). The image is stitched from 3 separate printing runs that each populate a subregion of the total array. The print rate for the array is 1.5 Hz, leading to an elapsed printing time of ~2 hours for the entire array. This demonstrates that the number of experiments that can be constructed on a single substrate using PDM (10^4^) is two orders of magnitude larger than the typical scales used in well plate fluid handling (10^2^).

### On-demand printing of single cells and complex cell combinations

Biological systems comprise populations of heterogeneous cells that are important to the form and function of the system as a whole. While there are now good methods for high throughput analysis of single cells, fewer methods exist for constructing and studying cell combinations. This would be important, for example, for reconstituting cellular interactions *in vitro* to study the impact of cellular heterogeneity on tissue function. Additionally, combinations of cells could be used for studying tissue self-organization, immune-tumor interactions, or the effects of tumor heterogeneity on drug response. Methods based on random encapsulation in droplets or chambers do not allow generation of defined cell combinations (18).

PDM provides an elegant solution to generating any desired combination of cells, by deterministically and repeatedly printing single cells of different type to droplet arrays. To illustrate this, we use the instrument to generate defined combinations of green (calcein green) and red (calcein red) stained PC3 prostate cancer cells. The cells are introduced into the printer as a mixed suspension and encapsulated into droplets randomly. The cell concentrations are set so that most droplets are empty but 5% contain a single red or green cell. They are scanned for fluorescence immediately after generation, to determine if they contain a single cell, appearing as bright red or green droplets (Fig. 3a). Empty droplets appear as the dominant feature on the scatterplot at low fluorescence values, while droplets containing red and green cells appear double positive at the upper-right of the scatter plot. In addition to a single cell, each array location is printed with 3 buffer droplets, to bring it to a final volume of ~1 nL. To generate defined arrays of printed cells, we program the instrument with an appropriate file, generating 100 positions containing alternating single red and green cells, a subsection of which is shown in Fig. 3b. We confirm proper loading of the array by imaging and find that 98 contain the correct contents. Close inspection reveals that positions missing cells have fabrication defects, an artifact of the laser etching method used to pattern electrodes, that should be eliminated when more reliable lithographic wet etching methods are used. By comparison, randomly loading an array with a suspension that contains an average of 0.5 red and 0.5 green cells per each array position, should lead to 36.8% empty wells, 36.8% wells with single cells, and 26.4% of wells with 2 or more cells. In practice, single cells are usually loaded into microscale reactors at ≤ 0.1 cells / well limiting dilution, leading to > 90% empty reactors.

**Figure 3.**
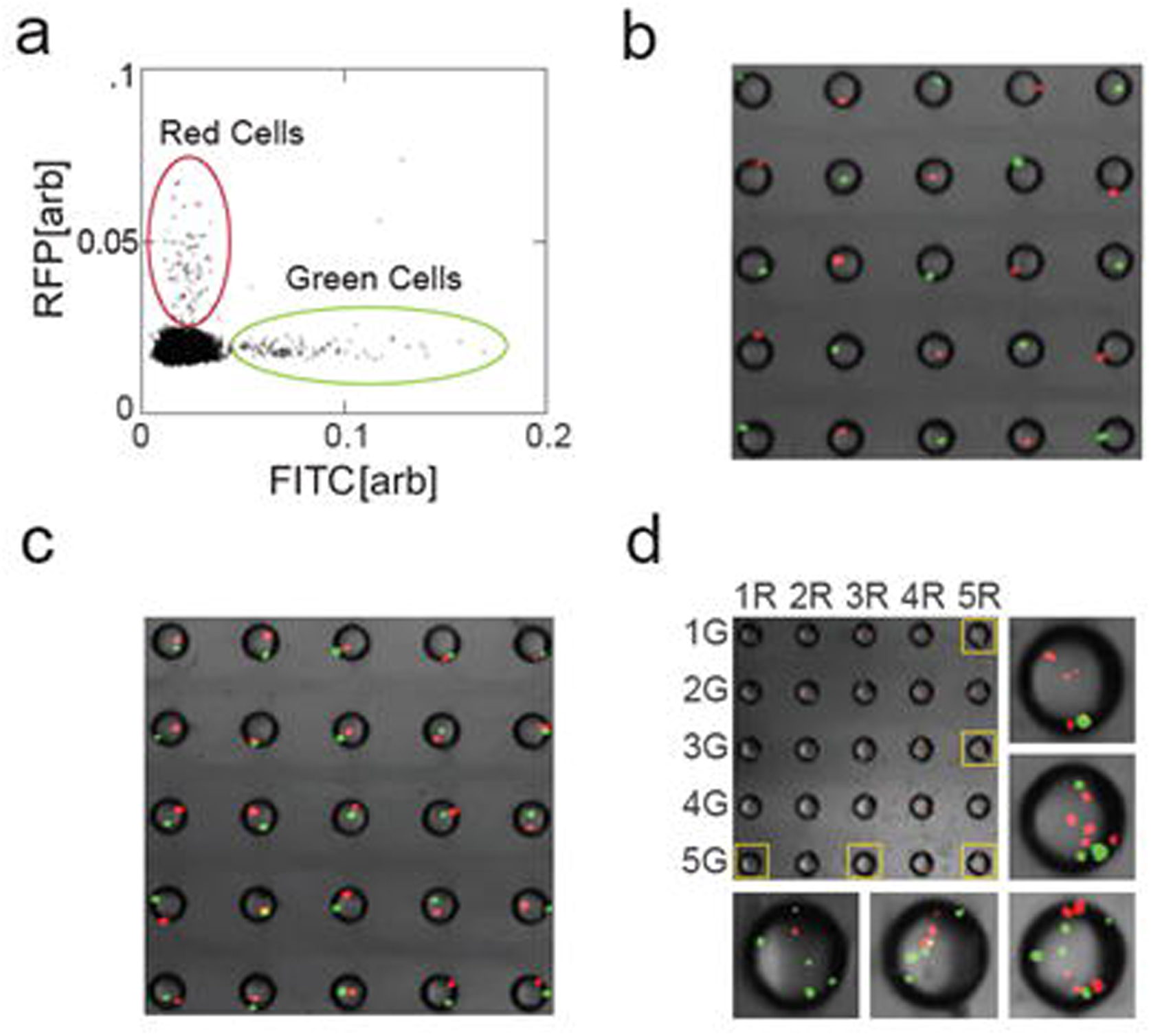
Cell printing. a) Flow cytom etry data for an em ulsion formed from a mix of calcein green- and calcein red-stained PC3 cells. b) 25-position array where every position is printed with alternating green- and red-stained single cells. c) 25-position array where every well is printed with both a single green- and red-stained cell. d) 25-position array printed with 1-5 green-stained cells on the vertical axis, and 1-5 red-stained cells on the horizontal axis.

In addition to the study of cell-cell interactions, numerous applications require the deterministic loading of defined numbers of cells or beads into droplets and wells, including new high throughput assays to detect secreted proteins for biologic drug screens (19) or to purify, barcode, and sequence the transcriptomes of thousands of single cells (20). The encapsulation of combinations of discrete entities like cells and beads, however, is usually inefficient, yielding a large majority of improperly loaded droplets. These droplets waste cells and reagents, and can contaminate putatively single cell data with data from multiple cells that otherwise appear indistinguishable.

PDM provides a robust way to deterministically generate any desired combination of cells and beads. The discrete entities are dyed to make them distinguishable and sorted-on-demand as needed to print the combinations, such as a red and green cell at every position (Fig. 3c). Of 100 printed positions, we again find 98 contain exactly one cell of each type. Randomly loading a nanowell array with a suspension containing an average of one cell of each type for every well would lead to 13.5% of the wells containing exactly one red and one green cell. Loading wells with two cell types at ≤ 0.1 cells / well limiting dilution, leads to > 99% of the wells containing undesired contents. This example illustrates the compounded advantage of deterministic cell delivery in constructing multicellular experiments.

Using PDM, arrays containing complicated combinations of cells are no more difficult to construct than single cell arrays and use exactly the same hardware: the print file is simply updated to specify the requisite printing instructions, and the printer runs them. This enables entirely new experiments that depend on multiple cell types. To illustrate the ease with which PDM can accomplish this otherwise impossible feat, we print complex combinations of cells: in which the number of cells per position systematically increases along an axis (Fig. 3d). This demonstrates that a large and reproducible variety of cellular consortia may be produced from a single mixed cell suspension.

### Single cell calcium release assay

Calcium signaling regulates diverse cellular processes such as cell motility, muscle contraction and gene transcription (21). Due to its diverse functionality, cellular Ca^2+^ level is highly regulated and it is maintained at low levels. The cellular Ca^2+^ level is regulated via cellular transmembrane channels as well as Ca^2+^ storage organelles. Intracellular Ca^2+^ release is a rapid event and monitoring intracellular Ca^2+^ changes could provide useful information regarding cellular response to a stimulus. To emphasize the versatility and sensitivity of experiments constructed using PDM, we utilized a fluorescence-based Ca^2+^ detection assay to demonstrate cellular response to stimuli, resulting in intracellular Ca^2+^ concentration changes. Here we show a high-throughput method for monitoring single cells in a time sensitive assay.

The demonstration application combines several key abilities of PDM: the ability to construct nanoliter volumes with variable reagent conditions, the controlled delivery single cells to these volumes, and the time dependent observation of precisely constructed microscale experiments. PC3 prostate cancer cells are used in the experiments, and are incubated with a green-fluorescing Ca^2+^ indicator dye. Intracellular calcium release is induced through the membrane depolarization caused by the titrated addition of potassium chloride (KCl). Bulk staining shows significant cell-to-cell variation both in the presence and absence of a stimulus (200 mM KCl) (Fig. 4a). Cells that were induced by 200 mM KCl show an average brightness that is 225% greater than non-stimulated cells. Using PDM, we can vary the stimulus and record single cell response with an arbitrary number of biological replicates. Single cell Ca^2+^ release experiments were constructed by printing each position in a 5 by 20 array with 0 - 200 mM KCl, then returning to each position and adding a droplet with a single cell treated with the Ca^2+^ indicator dye. Because Ca^2+^ signals are transient, imaging is performed in real time, lagging the single cell addition by 10 s. Due to the relatively dim signals from the Ca^2+^ indicator dye, autofluorescence of the epoxy used to construct the wells is visible. Wells that received higher concentrations of KCl are more likely to contain a cell with a detectable Ca^2+^ signal, and the cells contained within these wells are generally brighter. For wells titrated with 0, 50, 100, 150, and 200 mM KCl, there were 14, 12, 15, 19, and 19 wells with detectable cells, respectively. Comparing visible cells from 0 and 200 mM KCl single cell experiments, the induced cells have an average brightness that is 205% brighter than non-induced cells. The COVs for 200 mM KCl induced bulk and single cell experiments are 43% and 41%, respectively. The similarity in results between bulk and single cell assays demonstrate the ability of PDM to perform biologically relevant, single cell assays in miniaturized format.

**Figure 4.**
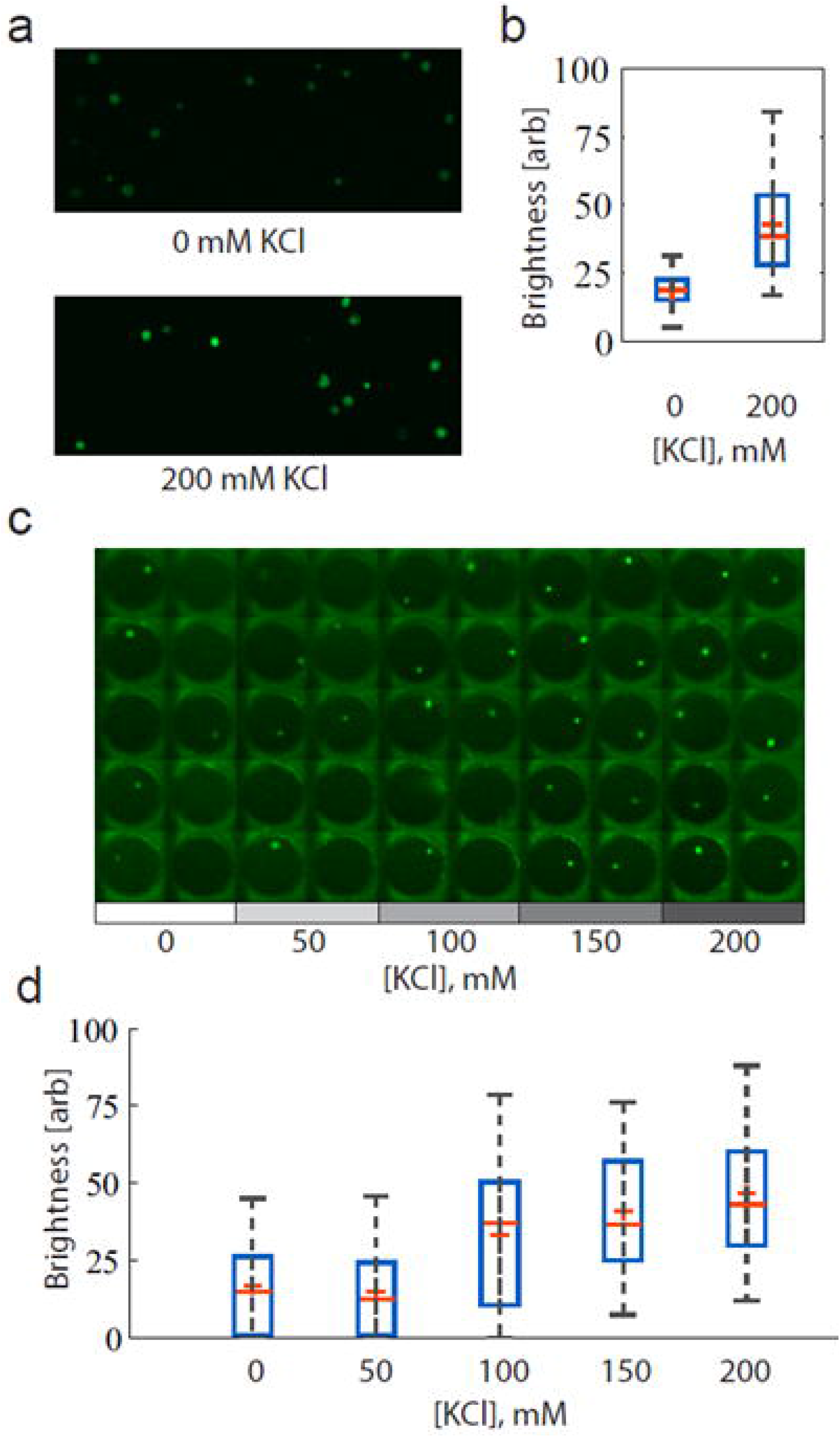
Intracellular calcium release assay. a) Assay results with and without the addition of 200 mM KC1 to induce Ca2+ release. b) Box plots of bulk cell assay data. Boxes represent 1st-3rd quartiles and whiskers give 5%-95% values. c) Assembled GFP channel imaging data from 50 of 100 array positions from single cell experiments. d) Box plots of single cell assay data. Boxes represent 1st-3rd quartiles and whiskers give 5%-95% values.

## Discussion

### Arrayed droplets are compatible with most measurement modalities

A common limitation in flowing droplet microfluidics is that the approach is only compatible with assays that provide bright signals and a high signal-to-noise ratio. Until now, this has restricted the approach to fluorescence or uncommonly sensitive absorbance assays, thus limiting the generality and usefulness of the technique. A critical advantage of PDM is that, as we show, the static droplets can be imaged and observed for long durations, which allows integration of weak signals into the detectable range. Imaging-based methods for absorbance and chemiluminescence have been demonstrated and should be readily applicable to printed droplets (22, 23). Similarly, optical spectrographic methods, like Raman spectroscopy, that are too weak to detect in flowing droplets, have been demonstrated using imaging and should adapt well to printed droplet arrays (24). More ambitiously, recent studies have performed electrospray ionization mass spectrometry on flowing droplets (25). The arrayed nature of printed droplets and ability to engineer the substrate to enable signal enhancement (26), provide unique opportunities for implementing it in an intelligent and useful way, especially in conjunction with laser ionization methods like MALDI, SAMDI, and NIMS that are compatible with the high throughputs required of future single cell biology applications (26–28). The ability to use a broader number of measurement techniques common to the biological sciences should greatly broaden the generality of droplet microfluidics.

### Arrayed droplets enable more controlled and intricate screens

An area into which droplet microfluidics has made significant headway is in the screening of enzymes via fluorescence-activated droplet sorting (1). However, a limitation of this approach is that sorting decisions must be made on an end-point measurement, because it is not possible to maintain droplets in registry as would be needed to perform time-dependent measurements. With PDM, time-dependent measurements are straightforward because each droplet is positioned at a single spot and can be imaged over time.

An additional benefit of arraying droplets is that the overlay fluid can be engineered to optimize for the chemistry of the particular screen. Since coalescence between arrayed droplets is not possible, surfactants that are the primary source of micelle-mediated leakage between droplets can be removed, reducing assay blur and enhancing measurement accuracy (29). Indeed, the oil can be replaced with another liquid or gaseous phase altogether, if necessary, to contain molecules that are natively soluble in fluorinated oil.

The spatial registration of droplets provides unique opportunities for maximizing the information and material recovered from a screen. For example, because printing is deterministic and programmable, the starting composition of every droplet is known, allowing straightforward relation of biochemical inputs to reaction outputs.

### Single cell multi-omics

We are just beginning the era of high throughput single cell analysis and its impact on our understanding of biology. But the complexity of biology is almost incomprehensible: In addition to characterizing large numbers of single cells, each cell should be characterized as deeply as possible, especially by combining many independent forms of single cell analysis. However, it is often challenging to integrate multiple analyses together, especially ones that are destructive. Additionally, at the conclusion of a screen, it is often desirable to recover live cells that can be propagated and studied further, or used to seed a new generation of cultures (30). With PDM, such single cell multi-omic measurements become feasible. Cells can grow to high densities in microfluidic droplets, and droplets can be split and printed to independent positions on the substrate. These subsamples provide multiple clonal colonies seeded from single cells that can be subjected to different analyses, including destructive ones. Even more, a pristine subsample can be preserved, providing live sister cells for recovery at the screen’s conclusion. Such workflows would be valuable in synthetic biology to evolve cells to produce molecules that can only be detected by destructive analyses, like mass spectrometry. Indeed, the majority of interesting molecules, including pharmaceuticals and chemical building blocks, fall into this category.

### Combinatorial biology

Biology is a science of combinations: The combinations of genes that give rise to a specific phenotype, chemical factors that induce differentiation into a specific state, and cells from which functional tissues and whole organisms emerge. Hence, it is a major limitation that, currently, we have few methods for performing high throughput combinatorial studies with cells. PDM allows this because it is possible to print arrays of droplets containing distinct, defined combinations of chemicals and cells. This should be valuable for screening chemical conditions that induce desired cell states, identifying combinations of microbes that grow under specific conditions, or combinations of cells that form functional tissue structures.

### Conclusion

We describe a new kind of microfluidics, PDM, that combines the flexibility and programmability of microliter dispensing with the scalability and single cell sensitivity of flowing droplet microfluidics. The core of this approach is a microfluidic sorter coupled to a motorized stage, used as a deterministic cell and droplet dispenser. In addition to allowing complex arrays of droplets and cells to be generated that are impossible by other means, PDM provides many other advantages, making it valuable for applications across subdisciplines of biology. Moreover, with the throughput increases that will result from easily implemented technical improvements, the approach can generate arrays of high complexity quickly and cost-effectively. This will allow new kinds of analyses on single cells and combinations of different cell types, integrating together multiple measurement modalities, both destructive and non-destructive, while permitting live cell recovery.

## Materials and Methods

### Printer operation

Printing reagents are loaded into syringes, attached to the print head using small bore polyethylene tubing (PE-2, SCI, Lake Havasu City, AZ), and mounted on syringe pumps (New Era, Farmingdale, NY). The unsorted waste outlet of the print head is coupled to a syringe pump running in reverse to ensure that roughly equal amounts of carrier oil exit the print nozzle and the waste channel. Print heads are mounted on an XYZ micromanipulator, so that the print nozzle can be brought in close contact with the printing substrate mounted on an epifluorescence microscope (AE31, Motic, Hong Kong). When the experiment is started, droplet detection data is acquired by a field programmable gate array (FPGA, National Instruments, Austin, TX) and displayed using a custom Labview application. The position of the print nozzle relative to nanowells can be viewed from below through the transparent substrate using a 4X or 10X objective. The relative position of the print nozzle to the printing substrate is achieved through the automated control of a mechanical stage (MS-2000, ASI, Eugene, OR). Sorting gates can be set within the application to trigger the FPGA to send a signal to the sorting electrode on the print head via a high voltage amplifier (609E-6, Trek). Print runs are implemented by uploading a print file, which give a sequential list of droplets to be delivered to each position on the substrate array.

